# Hash-based core genome multi-locus sequencing typing for *Clostridium difficile*

**DOI:** 10.1101/686212

**Authors:** David W Eyre, Tim EA Peto, Derrick W Crook, A Sarah Walker, Mark H Wilcox

**Author notes:** Profs Walker and Wilcox contributed equally. Corresponding author: David Eyre.

## Abstract

**Background:** Pathogen whole-genome sequencing has huge potential as a tool to better understand infection transmission. However, rapidly identifying closely-related genomes among a background of thousands of other genomes is challenging.

**Methods:** We describe a refinement to core-genome multi-locus sequence typing (cgMLST) where alleles at each gene are reproducibly converted to a unique hash, or short string of letters (hash-cgMLST). This avoids the resource-intensive need for a single centralised database of sequentially-numbered alleles. We test the reproducibility and discriminatory power of cgMLST/hash-cgMLST compared to mapping-based approaches in *Clostridium difficile* using repeated sequencing of the same isolates (replicates) and data from consecutive infection isolates from six English hospitals.

**Results:** Hash-cgMLST provided the same results as standard cgMLST with minimal performance penalty. Comparing 272 pairs of replicate sequences, using reference-based mapping there were 0, 1 or 2 SNPs between 262(96%), 5(2%) and 1(<1%) pairs respectively. Using hash-cgMLST or standard cgMLST, 197(72%) replicate pairs had zero gene differences, 37(14%), 8(3%) and 30(11%) pairs had 1, 2 and >2 differences respectively. False gene differences were clustered in specific genes and associated with fragmented assemblies. Considering 413 pairs of infections within ≤2 SNPS, i.e. consistent with recent transmission, 266(64%) had ≤2 gene differences and 50(12%) ≥5 differences. Comparing a genome to 100,000 others took <1 minute using hash-cgMLST.

**Conclusion:** Hash-cgMLST is an effective surveillance tool that can rapidly identify clusters of related genomes. However, cgMLST/hash-cgMLST generates potentially more false variants than mapping-based analysis. Refined mapping-based variant calling is likely required to precisely define close genetic relationships.

## Introduction

The rapid development of pathogen whole-genome sequencing offers huge potential for better understanding the epidemiology of many infections. When trying to intervene to stop transmission, it is often important to identify the most closely genetically-related organisms already sequenced, as these represent potential recent sources of infection or cases that share a common infection source. However, the rapidly growing scale of data generated makes identifying these closely-related genomes among a background of many thousands of other genomes very challenging.

Three main approaches can be taken to identify closely-related genomes. Comparing single nucleotide polymorphisms (SNPs) identified following mapping to a reference genome offers high precision, e.g.^1^ but, despite efforts to optimise computational approaches^2^, is relatively slow. In contrast, k-mer based approaches based on hash algorithms, e.g. MASH^3^, are fast, but the inherent dimensionality reduction (e.g. summarising the whole genome as 500 hash strings) makes these approaches less well suited to fine-scale transmission analyses. Core genome multi-locus sequencing typing (cgMLST)^4^ potentially provides a solution; genomes are summarised as a list of ∼2000-3000 numbers, with each number representing the unique sequence of each core gene. This summary enables more rapid comparisons as, taking the example of *Clostridium difficile*, only 2270 gene allele numbers need be compared,^5^ rather than having to compare 4.3 million base pairs of sequence data for SNPs. A drawback of cgMLST as described to date is that it requires a centralised database of alleles of each gene to be maintained, so that cgMLST profiles generated by different laboratories are comparable. This centralised support can potentially be provided by academic, public health or commercial organisations, but any given scheme’s sustainability is potentially limited by the funding available to support it. Additionally, for some pathogens, including *C. difficile*, several competing cgMLST/whole-genome-MLST schemes (e.g. Enterobase [University of Warwick, UK], cgmlst.org [Ridom GmbH, Germany] and BioNumerics [BioMérieux, France]) containing different genes and profiles have been developed; the latter two being associated with a commercial platform for processing sequencing data.

We therefore propose an alternative to cgMLST as described to date. Instead of maintaining a database of alleles, each allele is reproducibly converted to a unique hash, or short string of letters. This compresses each item of identical data to the same smaller representation based on the sequence of an allele alone. Therefore, this process can be undertaken independently in different laboratories without the need to maintain or subscribe to a central database, but still generates summary data in a reproducible form that can be exchanged by laboratories. This distributed approach avoids the potentially costly need maintain a central database.

This study has two main aims. Firstly, to demonstrate an implementation of hash-based cgMLST, and to test whether hash-cgMLST profiles can be compared without a significant performance penalty compared to standard cgMLST; and secondly to test the reproducibility and discriminatory power of cgMLST compared to SNP-based typing. The discriminatory power of cgMLST has been previously explored, e.g.^5–8^, however how cgMLST gene differences relate to SNP distances has not been comprehensively assessed. Instead it is postulated that small numbers of SNPs are likely to fall in different genes, and so SNP distances and gene differences are likely to be similar for closely related isolates. We evaluate the extent to which this assumption holds. Related to this, only limited assessments of the reproducibility of cgMLST have been undertaken. The largest study to date involved the same *Staphylococcus aureus* DNA from 20 isolates undergoing sequencing in 5 laboratories.^9^ In this setting, in 80 comparisons (i.e. 20 sequences from 4 laboratories compared with the baseline laboratory) only 3 false gene differences were identified. We investigate whether these results can be replicated in *C. difficile*.

## Methods

### Hash-cgMLST

Using the cgMLST scheme of Bletz *et al*,^5^ the first allele for each of the 2270 genes was used to create a BLAST search query. Following previous descriptions,^5,9^ BLAST searches for each gene required a 90% identity match, a matched length ≥99% of the query length and the matched gene to be free from ambiguous characters or premature truncation. To avoid apparent truncated genes arising from misassembly we checked the number of stop codons in the gene sequence, and only retained matches with a single stop codon. To avoid truncation arising from contig breaks we ensured that BLAST matches included the start and end of the query sequence. Other BLAST search parameters were: “evalue=0.01, word_size=11, penalty=−1, reward=1, gapopen=5, gapextend=2”. The resulting genes were either matched to the database available at cgmlst.org, i.e. standard cgMLST, or hashed using an md5 algorithm to create a 32-character hexadecimal string. The resulting cgMLST and hash-cgMLST profiles were saved as json files, i.e. a format that could readily be exchanged between laboratories. Where no BLAST match was found for a gene in the scheme an empty value was recorded, and that gene excluded in pairwise comparisons.

### Sequence data

During whole-genome sequencing of *C. difficile* undertaken in Oxford and Leeds, UK we have routinely re-sequenced a subset of isolates as part of our internal quality assurance. We searched our database for isolates sequenced more than once (see Supplementary Table S1 for NCBI/EBI SRA numbers). For a subset of these replicate sequences, the same extracted DNA was used to generate both sequences; for the remainder it was not documented in our laboratory information management system whether the same DNA extract was re-sequenced, or whether a fresh DNA extract was made from the same frozen isolate. Paired-end sequence data for both types of replicate were generated using Illumina technology, including on various iterations of the HiSeq platform and the MiSeq platform, with read lengths varying from 100-150bp in the majority of sequences (two 50bp sequences were also included).

To compare the discriminatory power of hash-cgMLST compared to SNP-based typing we processed 973 genomes from a previously published study of consecutive *C. difficile* over one year in six English hospitals using our hash-cgMLST and SNP pipelines (using data available in NCBI BioProject PRJNA369188).^10^

### Bioinformatic processing

For hash-cgMLST typing, raw sequence data underwent adapter trimming and quality trimming using bbduk.sh from the bbMap package (version 38.32).^11^ Stringent quality trimming was applied following Mellmann *et al*,^9^ both the left and right ends of each read were trimmed to a Q30 threshold (using bbduk parameters: “ktrim=r k=23 mink=11 hdist=1 tpe tbo qtrim=rl trimq=30”). Following this the number of bases remaining in the trimmed reads was divided by the length of the 630 reference genome^12^ (4290252 bp) to provide the mean high quality coverage, this was required to be ≥50 for a sequence to be included in the study. Appropriate quality trimming and adapter removal was confirmed using FastQC.^13^ To check for contamination with non-*C. difficile* DNA, the species origin of sequence reads was classified using Kraken2^14^ using the MiniKraken2_v1 database (built from the refseq bacteria, archaea, and viral libraries).

Reads were *de novo* assembled using SPAdes (version 3.11.1)^15^, with the “--careful” flag to reduce misassembly by using bwa-based mapping to confirm variants. Assembly quality metrics were obtained using the stats.sh script from bbmap. Following Bletz *et al*^5^ samples with assembly sizes (base pairs in contigs) >10% above or below the median size were rejected.

Reads (without stringent quality trimming) were also mapped to the 630 reference genome as described previously,^1,10,16^ using stampy^17^ for mapping and mpileup^18^ for variant calling, followed by quality filtering of variants. Variant calls were required to have a quality score of ≥30, be homozygous under a diploid model, be supported by ≥5 high quality reads including ≥1 read in each direction and a consensus of ≥90% of bases and not be within a repetitive region of the genome. See https://github.com/oxfordmmm/CompassCompact for example implementation. For inclusion, ≥70% of the reference genome needed to be called in the consensus sequence.

The bioinformatic pipelines used in this study for assembly and hash-cgMLST were written as NextFlow workflows^19^ and can be found at https://github.com/davideyre/hash-cgmlst.

### Analysis

Sequences meeting all quality thresholds (high-quality average coverage, assembly size, proportion of reference genome called) were compared. For replicate sequences, when an isolate had been sequenced more than twice, a random sequence was chosen as the baseline sequence with which all other sequences from the same isolate were compared, in order to avoid multiple counting.

Pairwise observed SNP differences between replicates and recombination-corrected SNP differences between other *C. difficile* genomes were obtained using Python scripts, PhyML^20^ and ClonalFrameML^21^ as previously described^10^ (https://github.com/davideyre/runListCompare). The number of cgMLST loci differences and number loci compared were obtained using Python (https://github.com/davideyre/hash-cgmlst).

## Results

Hash-cgMLST provided the same results as standard cgMLST with minimal performance penalty. Results are presented throughout using pairwise core-gene differences generated with hash-cgMLST as these were identical to standard cgMLST gene differences if novel alleles were accounted for.

### Comparison of hash-cgMLST and SNP typing performance in replicate sequences

A total of 374 sequences from 104 isolates passed all quality checks and were available for comparison to investigate the reproducibility of cgMLST for *C. difficile* transmission analyses. A median (interquartile range) [range] of 2 (2-3) [2-27] sequences were available per isolate. Comparing replicate sequences with a randomly selected baseline sequence for each isolate yielded 272 comparisons for analysis.

With perfect sequencing no variants would be expected between pairs of sequences from the same isolate (replicate pairs). Using reference-based mapping and variant calling there were 0 SNPs between 262 (96%) replicate pairs, 1 SNP between 5 (2%) pairs and 2 SNPs between 1 (<1%) pair, i.e. a mean 0.026 SNPs per pair which equates to 1 false SNP call per 39 sequences (Figure 1A). Based on the rate of *C. difficile* evolution and the extent of within host genetic diversity ≤2 SNPs are expected between >95% of cases related by recent transmission;^1^ therefore it is unlikely that transmission would be falsely excluded on the basis of the error rates seen.

**Figure 1.**
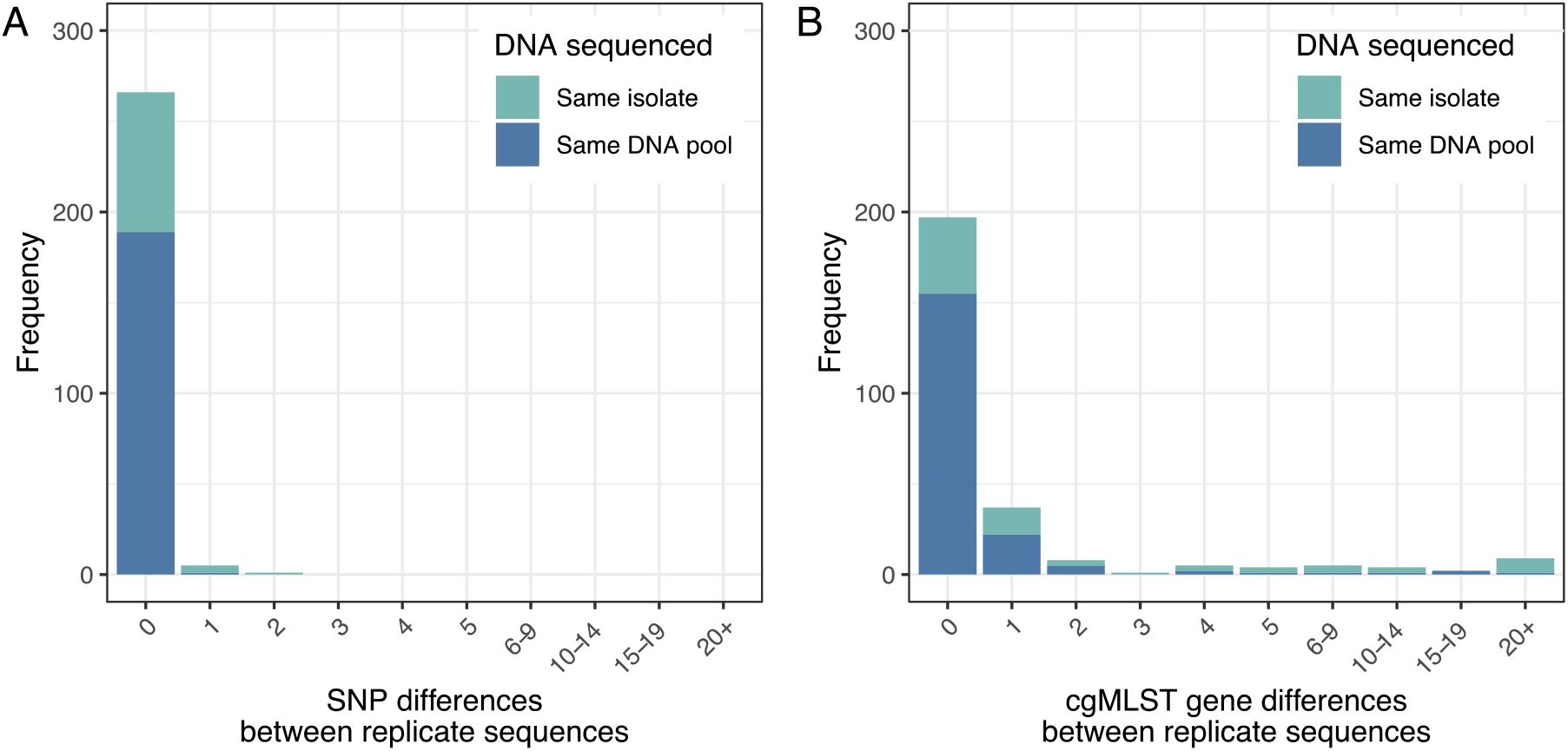
Observed differences using SNP typing (panel A) and hash-cgMLST (panel B) in 272 replicate sequence pairs. With perfect sequencing no variants would be expected between pairs of sequences from the same isolate. Pairs of sequences known to have been obtained from the same pool of DNA are shown in dark blue. Where information was unavailable on whether the same pool of DNA was used or a fresh DNA extract was made from the same isolate, this is shown in light blue.

Using either hash-cgMLST or standard cgMLST, 197 (72%) replicates pairs had zero gene differences, 37 (14%) pairs 1 difference, 8 (3%) pairs 2 differences, and 30 (11%) pairs had >2 differences, with a mean of 1.58 false gene differences per genome (Figure 1B) (test for symmetry considering 0, 1, 2, >2 SNPs or gene differences, p<0.001). Applying a threshold of >2 gene differences to rule out transmission (by analogy with SNP-based metrics^1,5^), the observed error rate would result in 11.0% (95% binomial confidence interval, CI, 7.6-15.4%) of transmission pairs being falsely excluded.

Restricting to the subset of sequences where sequencing was known to have been undertaken from the same pool of extracted DNA produced fewer false errors (Figure 1). Of 190 pairs, 189 (>99%) had 0 SNPs and 1 (<1%) pair had 1 SNP. From cgMLST, 155 (82%) pairs had 0 gene differences, 22 (12%) had 1 difference, 5 (3%) had 2 differences, and 8 (4%) had >2 differences. This would equate to 4.2% (95%CI 1.8-8.1%) of potential transmission pairs being falsely excluded by sequencing error using cgMLST with a threshold of >2 gene differences.

### Predictors of false cgMLST gene differences

The observation of substantially greater differences between replicates restricting to variation in the 2270 core genes versus considering variation across the whole genome is potentially counter-intuitive. However, it should be remembered that the whole-genome SNP approach depends on a different bioinformatic approach with sophisticated per variant quality filtering, whereas the cgMLST is based on *de novo* assembly with more limited quality filtering. We therefore investigated potential predictors of false cgMLST gene differences using the hash-cgMLST algorithm (which were identical to the standard cgMLST approach) to see if filtering could be improved. Although we had already restricted our analysis to only include sequences with a mean genome coverage of >50, we investigated whether a more stringent threshold would improve performance (Figure 2). There was marginal evidence that increased coverage was weakly associated with fewer cgMLST gene differences (Spearman’s rho −0.09, p=0.15), however >2 gene differences still occurred in pairs of samples where both had coverage of >150-fold. There were only 2 sequences in the dataset with 50bp reads, the remainder had 100 or 150bp reads. 24/222 (11%) sequence pairs where the minimum sequence length was 100bp contained >2 gene differences, compared to 6/48 (13%) in pairs with both 150bp reads (exact p=0.80).

**Figure 2.**
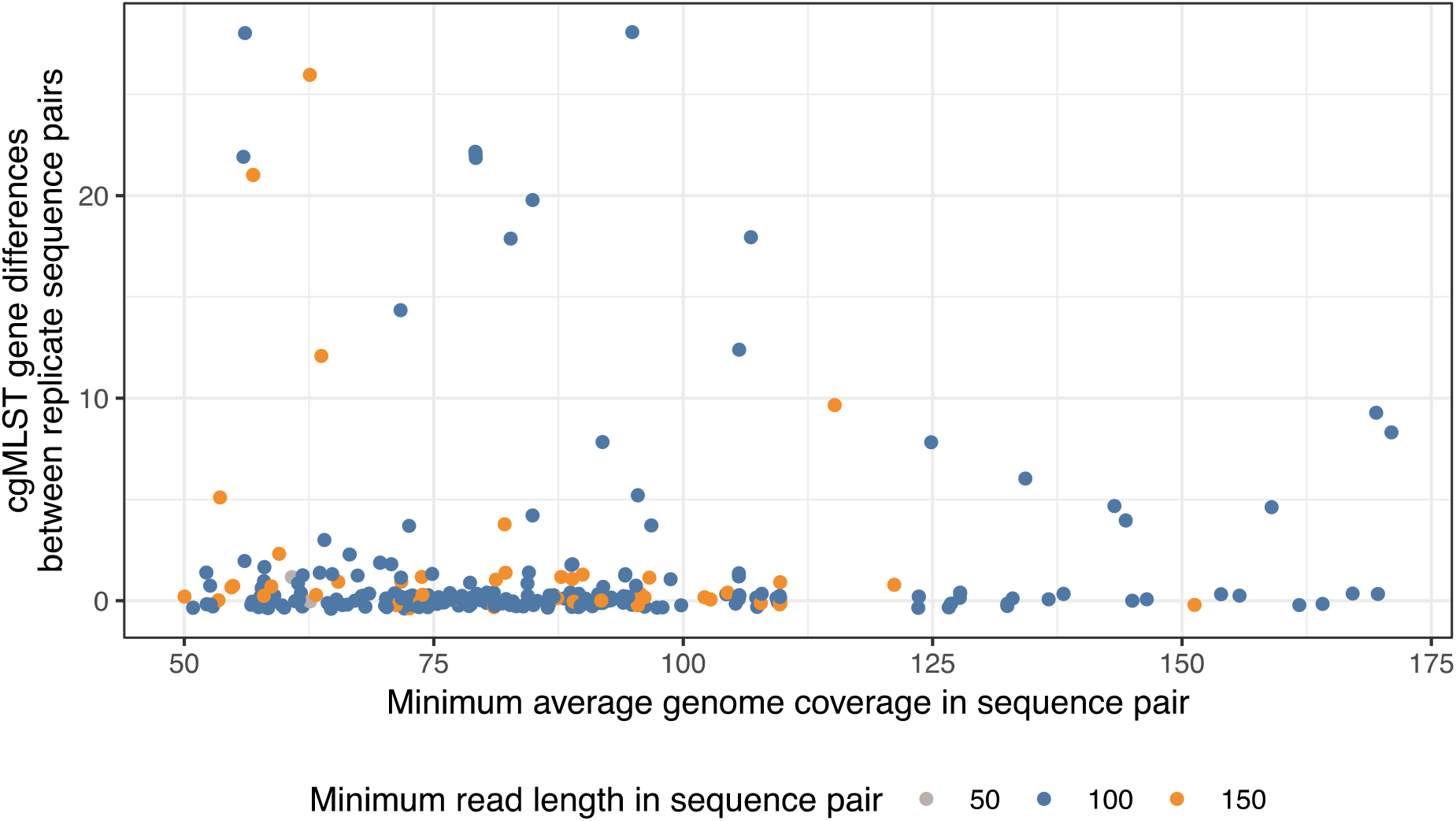
Relationship between hash-cgMLST gene differences in replicate sequence pairs and average genome coverage and read length. Jitter applied to points to assist visualisation.

The relationship between cgMLST gene differences and *de novo* assembly quality metrics is shown in Figure 3. Given the filtering applied, there was still an association between the number of false gene differences and the maximum absolute percentage deviation from the overall median assembly size (4165590bp) within each replicate pair (which was constrained to be ≤10% for inclusion in the analysis) (Spearman’s rho 0.22, p<0.001, Figure 3A, with both small and large assemblies contributing to this effect). L50 describes the minimum number of contigs required to achieve 50% of the assembly size, with higher values representing more fragmented lower quality assemblies. Higher values of L50 were associated with greater rates of false gene differences (Spearman’s rho 0.56, p<0.001). 15 (6%) of 257 pairs with both L50 values ≤125 had >2 false gene differences compared to 15/15 (100%) with one or more sequences with an L50 >125 (Figure 3B). Another measure of assembly fragmentation is the total number of contigs; higher numbers of contigs were also associated with greater false gene differences (Spearman’s rho 0.39, p<0.001, Figure 3C).

**Figure 3.**
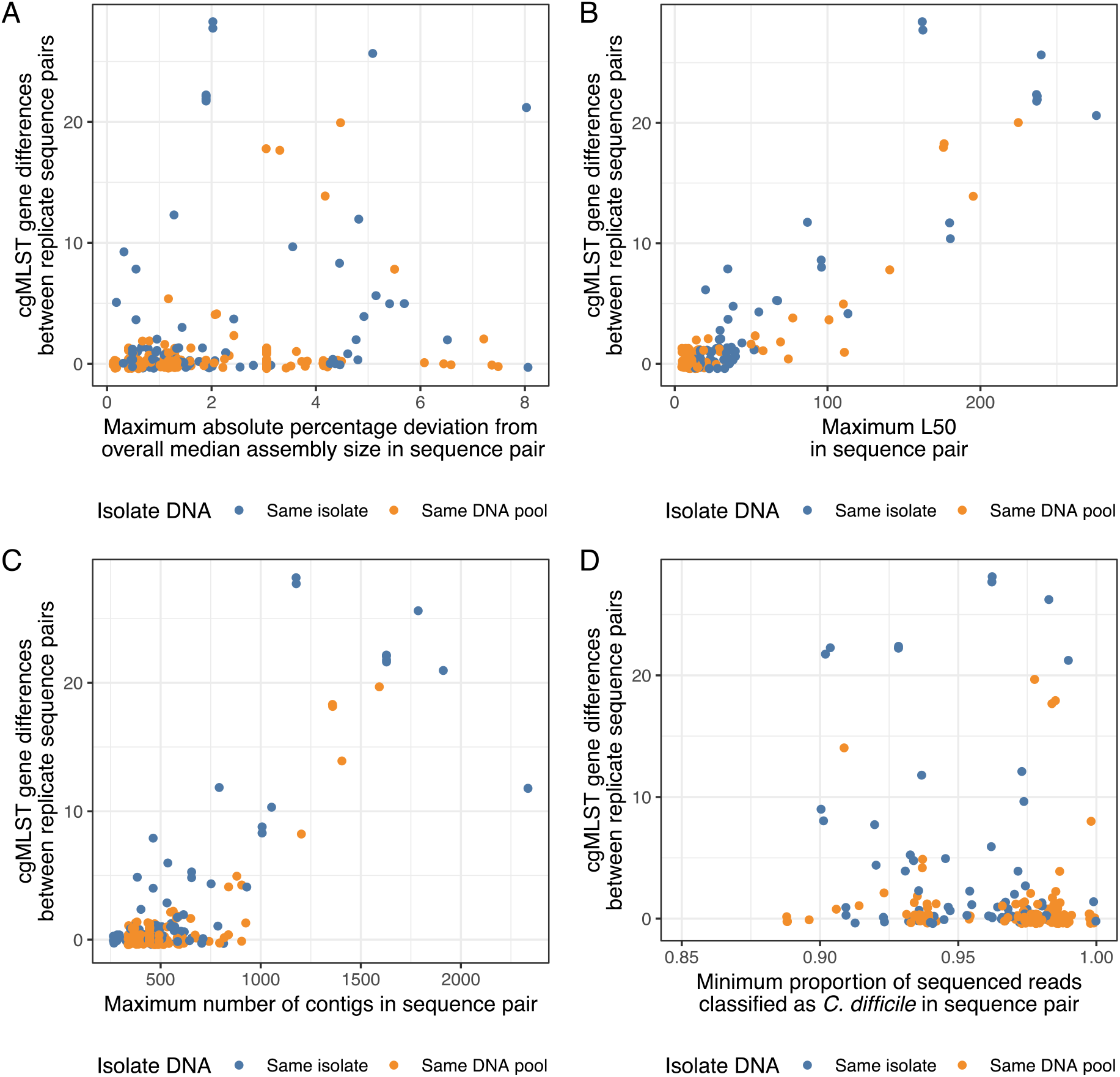
Relationship between hash-cgMLST gene differences in replicate sequence pairs and *de novo* assembly quality metrics and Kraken2 read classification. Jitter applied to points to assist visualisation. One point is omitted from Figure 3D for ease of visualisation with the proportion of reads classified as *C. difficile* of 0.64 and 1 gene difference.

Figure 3D shows the impact of the proportion of reads classified as *C. difficile* by Kraken2 on cgMLST gene differences. Within the dataset there was no evidence of significant contamination with a bacterial species other than *C. difficile* and the most common species was *C. difficile* in all samples. However, the proportion of reads that could not be classified at all varied from 0-11% between sequences with the exception of one replicate pair (36% and 24%). Higher rates of unclassified sequences were associated with higher false gene differences, but without any clear separation of the data on this basis (Spearman’s rho −0.23, p<0.001).

### Distribution of cgMLST gene differences in replicate sequences

The gene differences observed between replicate sequences disproportionately affected a small number of genes (Supplementary Table S2). Only 104 (5%) of 2270 genes contained differences within the replicate sequences. To avoid multiple counting, we evaluated the number of isolates that contained at least a pair of replicates with gene differences: gene CD630_16790 had differences affecting 58 (56%) (of the total 104) isolates, CD630_14380 differences in 40 isolates (38%), CD630_17540 in 21 (20%), CD630_24440 in 7 (7%), and 6 (6%) isolates had differences for CD630_13240, CD630_16000, CD630_32680, and 5 (5%) isolates had differences in CD630_20520, CD630_20910 and CD630_25780. The location of the differences observed was similar for a given gene across different isolates, compatible with consistent mis-assembly (Supplementary Table S2).

We investigated the impact of excluding genes with potentially higher rates of mis-assembly. If the 26 genes with gene differences affecting ≥3 isolates were excluded (Table S2), the number of the 272 replicate pairs with >2 gene differences reduced from 30 (11%) to 17 (6%). Restricting to the subset of 190 replicate pairs where sequencing was known to have been undertaken from the same pool of extracted DNA, the number of pairs with >2 gene differences reduced from 8 (4%) to 0 (0%) (Figure S1).

### Benchmarking

Samples were processed in parallel, with each sample using a single core from an Intel Xeon Gold 6150 2.70GHz 18-core CPU. For a single sample, the median (IQR) time to undertake quality control and read filtering was 3.6 (2.7-4.9) minutes and 27.4 (19.6-35.4) minutes to generate an assembly using Spades. From the assemblies creating a hash-cgMLST profile took 44.1 (43.5-44.9) seconds. Having made hash-cgMLST profile files, running on a single CPU core, to compare a single genome to 100,000 others took 40.4 seconds. In contrast 100,000 comparisons using a standard cgMLST approach took marginally less time, 38.7 seconds, after loading the profiles into memory.

### Comparison of hash-cgMLST and SNP typing in data from six English hospitals

We analysed 973 genomes from a previous study of *C. difficile* transmission in six English hospitals.^10^ Of these, 56 failed the assembly size threshold and 20 the coverage threshold (one also failing the assembly threshold), leaving 898 (92%) genomes for analysis. We considered all pairs of genomes within ≤2 SNPs and tested the extent to which the numbers of hash-cgMLST gene differences followed the number of SNPs (Figure 4A). Of 413 pairs of sequences within ≤2 SNPs, 266 (64%) were within ≤2 gene differences, 70 (17%) had 3 differences, 27 (7%) had 4 differences and 50 (12%) had ≥5 differences. Only 4 (1%) had >10 differences.

**Figure 4.**
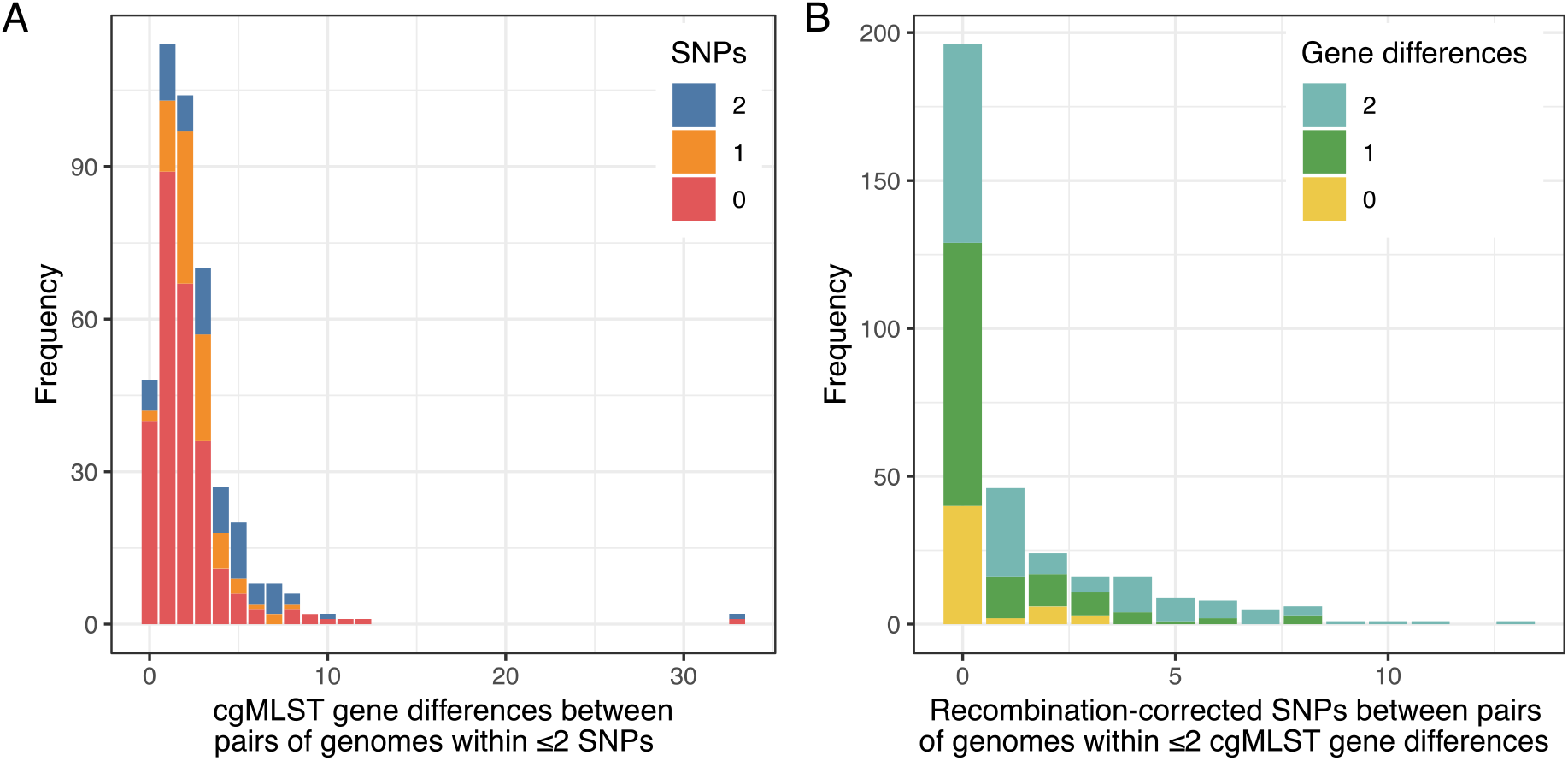
Relationship between hash-cgMLST gene differences and SNPS in *C. difficile* genomes from consecutive infections in six English hospitals. Panel A shows the distribution of hash-cgMLST gene differences between pairs of genomes within ≤2 SNPs. Panel B shows the distribution of SNPs within pairs of genomes within ≤2 gene differences.

Therefore, if a threshold of ≤5 gene differences was used to screen for isolates within ≤2 SNPs, the sensitivity of this approach was 93% (383/413), and the positive predictive value was 41% (383/932 pairs within ≤5 gene differences). Using a threshold of ≤10 the sensitivity and positive predictive value were 99% (409/413) and 13% (409/3046) respectively. Specificity was >99.9% with both thresholds (401791/401821 and 399703/399707 respectively). We also analysed the distribution of gene differences after excluding the 26 genes from the cgMLST scheme with potentially higher rates of mis-assembly identified from the replicate pairs. Using the modified scheme, if a threshold of ≤5 gene differences was used to screen for isolates within ≤2 SNPs, the sensitivity of this approach was 97% (402/413), and the positive predictive value was 25% (402/1600 pairs within ≤5 gene differences), with a threshold of ≤3 gene differences sensitivity was 96% (395/413) and positive predictive value was 46% (395/862) (Figure S2).

We also considered the distribution of SNPs within pairs of genomes with ≤2 gene differences using hash-cgMLST. Of 330 pairs of genomes, 266 (81%) were within ≤2 SNPs, with the maximum number of SNPs observed 13 (Figure 4B). Similar results were obtained excluding the 214 genomes with assemblies with an L50 value >125 (22% of all genomes).

## Discussion

Here we present the concept of hash-cgMLST as a tool for rapid comparison of bacterial sequencing data. This is a significant development over standard cgMLST approaches as it removes the need for a central database of alleles. Such databases require resource-intensive curation to ensure they are maintained to a high standard. Additionally, allele numbering is currently done consecutively in a single location, which is problematic with large datasets that span many laboratories; hashes also overcome this limitation. We also provide the code to run the algorithms developed.

This manuscript also highlights important limitations of common implementations of cgMLST as a tool for high resolution outbreak detection. Stringent filtering done on the basis of mapped data allows the number of false variant calls to be controlled; here we obtained around 1 false SNP for every 39 genomes sequenced. In contrast, fine-grained per base quality control is typically not implemented in studies using *de novo* assembly tools. As a result, higher rates of false variation were observed using cgMLST/hash-cgMLST, and were the main contributor to the counter-intuitive observation of more differences comparing 2270 genes than comparing the whole genome. It should be noted that undertaking SNP-based analyses from alignments of *de novo* assemblies without further filtering of variants would be similarly affected. These errors can be reduced by ensuring the assemblies studied are of high quality. Our data suggest that the previously described read quality trimming and filtering based on assembly sizes^5,9^ could be further improved by also only analysing samples with an L50 value of below ∼125. However, this stringent filtering would have resulted in 30% of a previously published dataset being unavailable for analysis, questioning its practicability.

We show that removing a small number of genes from the cgMLST scheme would likely improve performance, as a small subset of genes contained higher numbers of false gene differences (Table S2, Figure S1, Figure S2). This highlights the importance of assessing the performance of each cgMLST scheme created on a per species and scheme basis using appropriate test datasets which include replicate and closely-related sequences. As many of the errors seen appear to arise from mis-assembly, an alternative or complementary enhancement would be to improve upon the correction for misassembly already implemented in Spades, e.g. as in Enterobase^22^ where mapping to each assembly followed by filtering is used to assess the quality of the variants. It should be noted that the specific set of 26 genes excluded may in part be a property of the assembler used, and so care should be taken before extrapolating the current findings to other assemblers.

Our data also highlight that extrapolating the ≤2 SNP threshold for identifying genetically plausible transmission events to two (or three^5^) gene differences is likely to be inappropriate. Whether the additional variation we observed using hash-cgMLST (or standard cgMLST) is genuine or artefactual, much higher gene difference thresholds are required to identify samples within ≤2 SNPs. For public health applications optimised to identify potential transmission, using a threshold of ≤10 gene differences was 99% sensitive for identifying pairs of sequences within ≤2 SNPs, but resulted in around 6 genome pairs >2 recombination-corrected SNPs apart being identified for every 1 pair within ≤2 SNPs (positive predictive value 13%). In this scenario further SNP-based analysis based on mapping and filtered variant calling is likely to be required to determine which genomes are potentially related by recent transmission and which are not.

Hash-cgMLST allowed rapid comparison of many thousands of bacterial genomes within seconds, using a relatively unoptimized python script running on a single CPU core. As comparisons with other genomes can be easily divided into independent parts, this task is readily parallelisable. Using hash-cgMLST, it is therefore potentially possible to compare each new sequence generated with millions of previous sequences. The summaries of each genome produced, a roughly 130kb json file, are readily exchangeable between laboratories and could potentially be hosted alongside raw reads in sequence read archives. Although without further refinements hash-cgMLST may not allow high-precision fine-scaled transmission studies, it has the potential to dramatically reduce the search space for closely-related genomes, which can then be followed by more precise SNP-based analyses on a much smaller subset of genomes.

We observed a higher rate of ‘false’ gene differences between genomes where the sequences were potentially generated from separate DNA extractions of the same isolates, compared with genomes obtained from the same DNA extraction. It is therefore plausible that the differences observed represent true differences, but a form of variation that is much faster and more erratic than mutation rates based on filtered SNPs. The erratic nature of the variation observed is unlikely to be informative about recent transmission.

This study is potentially limited by not being an exhaustive investigation of all the potential options for filtering *de novo* assembly data, in particular further filtering of variants based on mapping reads back to assemblies may improve precision.^22^ Although we used Kraken2 to search for contamination with DNA from other species, contamination with *C. difficile* DNA from other samples processed concurrently may be an important contributor to some of the differences seen with hash-cgMLST, whereas resulting mixed calls can be filtered using mapped data.

In conclusion, appropriately quality controlled cgMLST can identify clusters of related genomes rapidly and is an appropriate tool for surveillance and reducing the search space in outbreaks. Refined variant calling based on mapping is likely required to precisely define close genetic relationships. Updates to the genes considered in the cgMLST scheme and to approaches for controlling for mis-assembly should be considered. This study highlights the need for detailed quality assurance to determine the performance of algorithms used for comparing genomes. Our hash-cgMLST implementation is freely available and provides an effective database-free approach to cgMLST.

